# TargetCall: Eliminating the Wasted Computation in Basecalling via Pre-Basecalling Filtering

**DOI:** 10.1101/2022.12.09.519749

**Authors:** Meryem Banu Cavlak, Gagandeep Singh, Mohammed Alser, Can Firtina, Joël Lindegger, Mohammad Sadrosadati, Nika Mansouri Ghiasi, Can Alkan, Onur Mutlu

## Abstract

Basecalling is an essential step in nanopore sequencing analysis where the raw signals of nanopore sequencers are converted into nucleotide sequences, i.e., reads. State-of-the-art basecallers employ complex deep learning models to achieve high basecalling accuracy. This makes basecalling computationally-inefficient and memory-hungry; bottlenecking the entire genome analysis pipeline. However, for many applications, the majority of reads do no match the reference genome of interest (i.e., target reference) and thus are discarded in later steps in the genomics pipeline, wasting the basecalling computation.

To overcome this issue, we propose TargetCall, the first fast and widely-applicable pre-basecalling filter to eliminate the wasted computation in basecalling. TargetCall’s key idea is to discard reads that will not match the target reference (i.e., off-target reads) prior to basecalling. TargetCall consists of two main components: (1) LightCall, a lightweight neural network basecaller that produces noisy reads; and (2) Similarity Check, which labels each of these noisy reads as on-target or off-target by matching them to the target reference. TargetCall filters out all off-target reads before basecalling; and the highly-accurate but slow basecalling is performed only on the raw signals whose noisy reads are labeled as on-target.

Our thorough experimental evaluations using both real and simulated data show that TargetCall 1) improves the end-to-end basecalling performance of the state-of-the-art basecaller by 3.31 × while maintaining high (98.88%) sensitivity in keeping on-target reads, 2) maintains high accuracy in downstream analysis, 3) precisely filters out up to 94.71% of off-target reads, and 4) achieves better performance, sensitivity, and generality compared to prior works. We freely open-source TargetCall to aid future research in pre-basecalling filtering at https://github.com/CMU-SAFARI/TargetCall.

## 1. Introduction

Genome sequencing, which determines the nucleotide sequence of an organism’s genome, plays a pivotal role in enabling many medical and scientific advancements [1–5]. Modern sequencing technologies produce increasingly large amounts of genomic data at low cost [6]. Leveraging this genomic data requires new fast, efficient, and accurate analysis tools.

Current sequencing machines are unable to determine an organism’s genome as a single contiguous sequence [7]. Instead, they extract and sequence subsequences of a genome, called *reads*. The length of the reads depends on the sequencing technology, and significantly affects the performance and accuracy of genome analysis. The use of *long reads* can provide higher accuracy and performance on many genome analysis steps [8–13].

Nanopore sequencing technology is one of the most prominent and widely-used long read sequencing technologies, for example used by Oxford Nanopore Technologies (ONT) sequencers [6, 14–18]. ONT sequencers rely on measuring the change in the electrical current when a nucleic acid molecule (DNA or RNA) passes through a pore of nanometer size [19]. The measurement of the electrical current, called a *raw signal*, is converted to a nucleotide sequence, called a *read*, with a step called *basecalling* [6, 7, 20–23]. Basecalling commonly uses computationally expensive deep neural network (DNN)-based architectures to achieve high basecalling accuracy [24,25]. However, the majority of this computation is wasted for genome sequencing applications that discard the majority of the basecalled reads. For example, in SARS-CoV-2 genome assembly, 96% of the total runtime is spent on basecalling, even though ≥99% of the basecalled reads are discarded later [26]. Therefore, it is important to eliminate wasted computation in basecalling.

Our **goal** in this work is to eliminate the wasted computation in basecalling while maintaining high accuracy and applicability to a wide range of genome sequencing applications. To this end, we propose TargetCall, the first widely applicable pre-basecalling filter. TargetCall is based on the key observation that typically the reason for discarding basecalled reads is that they do not match some *target reference* (e.g., a reference genome of interest) [26, 27]. We call these *off-target* reads. Our **key idea** is to filter out off-target reads before basecalling with a highly accurate and high-performance *pre-basecalling filter* to eliminate the wasted computation in basecalling of off-target reads.

Prior works [19,26,28,29] propose *targeted sequencing* techniques to discard off-target reads during sequencing to better utilize sequencers. These techniques can reduce wasted computation in basecalling and some could be partially repurposed for pre-basecalling filtering. However, all of them have at least one of three key shortcomings. First, some have low (77.5%-90.40%) sensitivity (i.e., they falsely reject many on-target reads) [19, 28]. Second, some are limited to a target reference size smaller than ~ 100 Mbp, i.e., they cannot be applied to longer target references, such as the human reference genome [19, 26, 28]. Third, some require re-training neural network based classifiers for each different application and target reference [29].

TargetCall consists of two main components: 1) *LightCall*, a light-weight basecaller with a simple neural network model that outputs noisy reads with high performance; and 2) *Similarity Check* to compute the similarity of the noisy read to the target reference. LightCall’s model is 33.31 × smaller than the state-of-the-art basecaller, Bonito’s model [30], which improves the basecalling speed substantially with small (4.85%) reduction in basecalling accuracy. Although LightCall’s basecalling is not accurate enough for some of the downstream analyses [31], it is sufficient for Similarity Check to perform pre-basecalling filtering. TargetCall overcomes all three limitations of prior methods. First, Similarity Check’s high sensitivity enables TargetCall’s sensitivity to be significantly higher than prior targeted sequencing approaches. Second, LightCall’s performance is independent of the target reference size, which enables TargetCall to be applicable to target reference sizes for which prior works were unapplicable. Third, unlike prior approaches that require re-training of the network for each application and target reference, LightCall does not need to be re-trained.

### Key Results

We evaluate the performance and accuracy impact of TargetCall on the state-of-the-art basecaller Bonito [30], and compare TargetCall with two state-of-the-art targeted sequencing methods, UNCALLED [28] and Sigmap [19], repurposed as pre-basecalling filters. TargetCall 1) improves the end-to-end performance of basecalling by 3.31 × over Bonito, 2) precisely filters out 94.71% of the off-target reads, and 3) maintains high sensitivity in keeping on-target reads with 98.88% recall. We show that TargetCall provides near-perfect downstream analysis accuracy even after losing 1.12% of on-target reads. We demonstrate that TargetCall improves performance by 1.95×/3.07× over prior works UNCALLED/Sigmap, respectively, on the *subset* of use cases that these tools are applicable to.

This paper makes the following contributions:

- We introduce the first widely applicable pre-basecalling filter that eliminates the wasted computation in basecalling by leveraging the fact that the majority of reads are discarded after basecalling.
- We propose LightCall, a light-weight neural network model that significantly increases the performance of basecalling with minor reductions in basecalling accuracy.
- TargetCall provides larger performance and accuracy benefits compared two state-of-the-art targeted sequencing works for basecalling while being applicable to large target reference sizes as well.
- To aid research and reproducibility, we freely open source our implementation of TargetCall.

## 2. Background & Motivation

### 2.1. Genome Sequence Analysis

Nanopore sequencing is a widely used sequencing technology that produces long reads with high-throughput [6, 14–18]. Oxford Nanopore Technologies (ONT) is a company that produces sequencers that perform nanopore sequencing. The ONT sequencers output the raw electrical signal as a representation of the nucleic acid molecule that passes through the nanopore [19]. The sequenced data is analyzed following the three key steps: basecalling, read mapping, and variant calling [32]. Basecalling is the process of translating this raw signal into a sequence of nucleobases (A,C,G,T) to generate the reads. Basecalling is a nontrivial task due to the high noise levels of raw signals produced by sequencers. State-of-the-art basecallers (e.g., Bonito [30]) use DNN-based architectures to provide high accuracy for this non-trivial task [30, 33–39]. Due to computationally-inefficient and memory-hungry DNN architectures used, basecalling is the most time consuming step in genome sequence analysis. As an example, a real system study [40] shows that basecalling consumes 84.2% of total execution time. The second step, read mapping, computes the sequence alignment of the read to the reference genome (i.e., representative genome sequence of a species) [1, 41–43]. The third step of the pipeline is variant calling that computes variants of the genome with respect to the reference genome using aligned reads [44]. The same study shows that read mapping and variant calling steps consume 13.6% and 2.2% of total execution time respectively [40]. We conclude that basecalling is a major bottleneck of the genome sequence analysis pipeline, and it is critical to reduce basecalling execution time to improve performance of genome analysis.

### 2.2. Targeted Sequencing

Targeted sequencing is a method to selectively sequence reads from the reference genome of interest (i.e., target reference) during sequencing time [28]. ONT devices have the potential to enable computational targeted sequencing without the need for library preparation with a feature, known as *Read Until* [28,45]. ONT sequencers that support Read Until can selectively remove a read from the nanopore while the read is being sequenced. Read Until can be used to remove off-target reads during sequencing, eliminating wasted computation on off-target in basecalling. All targeted sequencing approaches work by labeling a read as on-target or off-target and stopping the sequencing of off-target reads immediately after labeling using Read Until. We discuss three groups of works that perform Read Until, classified based on their methodology to label the read. The first group converts the target reference into a reference raw signal and performs raw signal-level alignment [19,26,45]. The second group generates noisy sequence representations of the raw signal to compare them with the target reference [28, 46]. The third group of works utilize neural network classifiers to label the sequences [29,47]. All these approaches are explained in detail in Section 7. To our knowledge, none of these works can be fully repurposed as widely-applicable pre-basecalling filters to eliminate the wasted computation in basecalling for a wide range of genome applications.

#### Limitations of Targeted Sequencing Approaches

Even though these approaches can discard off-target reads from the genome analysis pipeline hence eliminating the wasted computation in basecalling, all have at least one of the following three key limitations. First, some [19, 28, 46] have low (77.5%- 90.40%) sensitivity which affects the accuracy of downstream analysis. These tools falsely reject a significant portion (~10%- ~23%) of the on-target reads. Second, some [19, 26, 28, 45] are not scalable to the long target references. This happens due to one of the following reasons: 1) use of signal-signal alignment algorithms whose complexity increases linearly with the target reference length [26, 45], 2) use of complex data structures to represent target reference in signal domain that are not scalable to long target reference lengths [19], and 3) use of probabilistic algorithms that scale poorly with the increasing target reference length [28]. These tools cannot be used for applications that require long target references, such as human reference. Third, some require neural network classifiers to be re-trained for each different application and target reference [29]. The re-training is required since the classifier is trained depending on the specific set of on-target and off-target reads. These approaches cannot be used in a wide range of applications without significant overheads.

## 3. Methods

Our goal in this work is to eliminate the wasted computation in basecalling using an accurate pre-basecalling filtering technique. To this end, we propose TargetCall, that can perform pre-basecalling filtering in *all* genome sequencing applications accurately and efficiently without any additional overhead. To our knowledge, TargetCall is the first pre-basecalling filter that is applicable to a wide range of use cases. TargetCall’s key idea is to quickly filter out off-target reads (i.e., reads that are dissimilar to the *target reference.)* before the basecalling step to eliminate the wasted computation in basecalling. We present the high level overview of TargetCall in Section 3.1, and explain its components in Sections 3.2 and 3.3.

### 3.1. High Level Overview

Figure 1 shows TargetCall’s workflow. First, TargetCall performs noisy basecalling on the raw signal using LightCall ❶. The output sequence of LightCall is a highly accurate but noisy read. Second, Similarity Check compares the noisy read of LightCall to the target reference for labelling the read as an on-target or off-target read ❷. TargetCall stops the analysis of off-target reads by removing them from the pipeline ❸, whereas the analysis of the on-target reads continues with basecalling following the usual genomics pipeline to maintain basecalling accuracy ❹.

**Figure 1:**
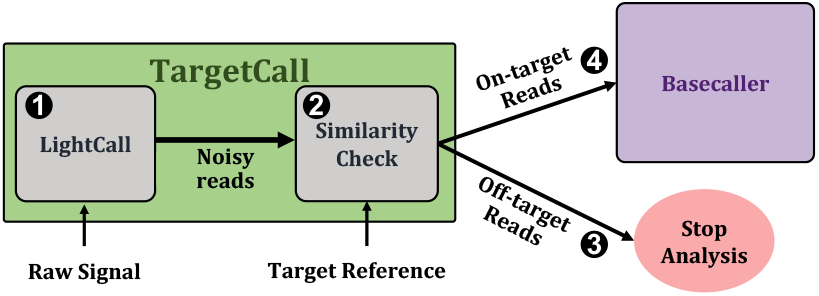
High-level overview of TargetCall.

The choice of the target reference depends on the specific genome sequencing application. The size of target reference is a major constraint in most prior works, limiting the generality of the prior approaches. We design TargetCall such that its performance is independent of the size of the target reference so that it is applicable to any genome sequencing application.

### 3.2. LightCall

LightCall, the first component of TargetCall, is a light-weight neural network based basecaller that produces noisy reads. The basecalling accuracy of LightCall is less than the state-of-the-art basecallers, however, it is still high enough for Similarity Check to determine if the read is an on-target read with respect to the target reference. We develop LightCall by modifying the state-of-the-art basecaller Bonito’s architecture in three ways: 1) reducing the channel sizes of convolution layers, 2) removing the skip connections, and 3) reducing the number of basic computation blocks. Prior work [48] shows that Bonito’s model is over provisioned, and we can maintain very high accuracy with reduced model sizes. We leverage this finding in TargetCall to reduce the channel sizes of convolution layers and number of basic computation blocks of LightCall.

Figure 2 shows the architecture of LightCall. Each block consists of grouped 1-dimensional convolution and pointwise 1-dimensional convolution. The convolution operation is followed by batch normalization (Batch Norm) [49] and a rectified linear unit (ReLU) [50] activation function. The final output is passed through a connectionist temporal classification (CTC) [51] layer to produce the decoded sequence of nucleotides (A, C, G, T). CTC is used to provide the correct alignment between the input and the output sequence. Our LightCall architecture is composed of 18 convolution blocks containing ~292 thousand model parameters (~33.35 × lower parameters than Bonito).

**Figure 2:**
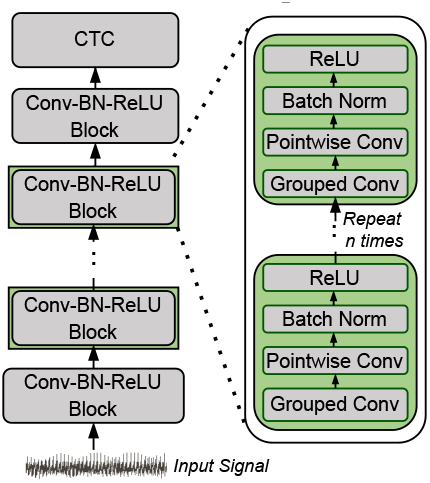
Overview of LightCall architecture.

Modern deep learning-based basecallers [20, 21, 36, 38, 39, 48, 52–54] incorporate skip connections to help mitigate the vanishing gradient and saturation problems [55]. Removing skip connections has a higher impact on basecalling accuracy. However, adding skip connections introduces the following three issues for performance. First, skip connections increases the data lifetime. The layers whose activations are reused in subsequent layers must wait for this activation reuse (or buffer the activations in memory) before accepting new input and continuing to compute. This leads to high resource and storage requirements due to data duplication. Second, skip connections introduce irregularity in neural network architecture as these connections span across non-adjacent layers. Third, skip connections require additional computation to adjust the channel size to match the channel size at the non-consecutive layer’s input. Therefore, we remove the skip connections as we can tolerate lower accuracy of LightCall to improve performance of LightCall.

LightCall splits a long read in raw signal format (e.g., millions of samples) into multiple smaller chunks (e.g., thousands of samples per chunk) and basecalls these chunks. LightCall uses the input signal to predict nucleotides as labels. The CTC layer assigns a probability for all possible labels at each timestep. The nucleotide with the highest probability is selected as the final output.

### 3.3. Similarity Check

After LightCall outputs the noisy read that approximately represents the raw signal, Similarity Check compares the noisy read to the target reference. For this task, we use a procedure common in many genome analysis, known as the sequence alignment. Sequence alignment computes the similarity between a read and a reference. We use the state-of-the-art read mapping tool that performs sequence alignment (i.e., minimap2 [56]) for Similarity Check. Similarity Check labels a read that is similar to the target reference as on-target. With the efficient index structure of minimap2, the performance of sequence alignment is almost independent of the length of the target reference genome, hence, TargetCall is scalable to large target reference genomes of Gbp (i.e., Giga base pair) length.

The accuracy of the computed sequence alignments is not high enough to represent the true alignment between the read and the target reference since the reads computed by LightCall are noisy. However, the labelling accuracy of Similarity Check for determining the on-target/off-target reads is high enough to provide sensitivity up to 99.45% in filtering reads. The minor inaccuracy of Similarity Check can be compensated with the high sequencing depth-of-coverage, the average number of reads that align to a genomic region, required for confident genome sequence analysis [57–60].

## 4. Use Cases of TargetCall

To show the applicability of TargetCall, we evaluate it on use cases with varying target reference sizes without compromising accuracy. We describe three use cases we use to evaluate TargetCall: (1) Covid Detection, (2) Sepsis Detection, and (3) Viral Detection. All three use cases contain a significant fraction of off-target reads that are eliminated using TargetCall to show the benefits of pre-basecalling filtering.

### Covid Detection

The first use case aims to filter reads coming from a small target reference to show TargetCall is applicable to use cases analyzed in targeted sequencing methods. We choose SARS-CoV-2 detection as a sample biological application where the goal is to detect the reads coming from a SARS-CoV-2 reference genome (~30 Kbp) from a sample taken from a human [26, 61].

### Sepsis Detection

The second use case aims to filter reads coming from a large target reference to show TargetCall is applicable to use cases where targeted sequencing methods are unapplicable due to their poor scalability with increasing target reference size. We choose sepsis detection [27, 62, 63] as a sample biological application where the goal is to delete the human reads from a human sample. Since the bacteria that is causing the disease is unknown, we cannot search for reads coming from a specific bacterial target. Instead, we apply TargetCall to filter-out reads similar to the target reference.

### Viral Detection

The third use case aims to filter reads from a collection of reference genomes to show TargetCall can correctly filter reads when the sample and the target reference have a high variety of species. This is to test the specificity of TargetCall in filtering reads when the on-target and off-target reads resemble each other more compared to previous use cases. We choose disease-causing viral read detection as a sample biological application where the goal is to detect the viral reads from a metagenomic sample of bacterial and viral reads [64].

## 5. Evaluation Methodology

### Evaluation System

Table 1 provides our system details. We use NVIDIA TITAN V to train and evaluate LightCall and baseline techniques. We use the state-of-the-art read mapper, minimap2 [56] for the Similarity Check module of TargetCall with -a and -x map-ont flags. The -a flag is used to compute sequence alignments and -x map-ont is used to configure minimap2 parameters for ONT data.

**Table 1:** System parameters and hardware configuration for the CPU and GPU.

|  |  |
| --- | --- |
| **CPU** | AMD EPYC 7742 [65]<br>@2.25GHz, 4-way SMT [66] |
| **Cache-Hierarchy** | 32×32 KiB L1-I/D, 512 KiB L2, 256 MiB L3 |
| **System Memory** | 4×32GiB RDIMM DDR4 2666 MHz [67] PCIe 4.0 ×128 |
| **OS details** | Ubuntu 21.04 Hirsute Hippo [68],<br>GNU Compiler Collection (GCC) version 10.3.0 [69] |
| **GPU** | NVIDIA TITAN V [70] 5120 CUDA Cores@1.2GHz, 12GiB HBM2<br>NVIDIA System Management Interface (NVIDIA-SMI) version 510.47.03 [71]<br>NVIDIA CUDA Compiler Driver (NVCC) version 11.1.105 [72] |

### Training Setting

We use the publicly available ONT dataset [30] sequenced using MinION Flow Cell (R9.4.1) [73] for the training and validation. The neural network weights are updated using Adam [74] optimizer with a learning rate of 2*e*^−3^, a beta value of 0.999, a weight decay of 0.01, and an epsilon of 1*e*^−8^.

### Baseline Techniques

We evaluate TargetCall’s performance and accuracy as a pre-basecalling filter by integrating it as a pre-basecalling filter to Bonito [30], which is the official basecalling tool developed by ONT. In Section 6.6, we evaluate two state-of-the-art targeted sequencing methods, UNCALLED [28] and Sigmap [19] repurposed as pre-basecalling filters to compare against TargetCall. We do not evaluate TargetCall against other targeted sequencing methods such as SquiggleNet [29] that cannot be trivially repurposed as pre-basecalling filters.

### LightCall Configurations Evaluated

To determine the final architecture of TargetCall, we test 5 different LightCall configurations. Table 2 lists the LightCall configurations evaluated.

The number of parameters each configuration has varies and affects the performance and accuracy of the model. The higher number of model parameters results in slower but more accurate models. We evaluate these 5 different LightCall configurations to explore the performance-accuracy trade-off in LightCall, and select the LightCall architecture that provides the best trade-off.

**Table 2:** Different LightCall configurations.

| Model Name | Basecalling Accuracy | Number of Parameters | Model Size (MB) |
| --- | --- | --- | --- |
| Bonito | 94.60% | 9,739K | 37.14 |
| $LC_{Main \times 2}$ | 90.91% | 565K | 2.16 |
| $LC_{Main}$ | 89.75% | 292K | 1.11 |
| $LC_{Main/2}$ | 86.83% | 146K | 0.55 |
| $LC_{Main/4}$ | 80.82% | 52K | 0.19 |
| $LC_{Main/8}$ | 70.42% | 21K | 0.07 |

### Evaluated Datasets

We use four real and one simulated dataset to evaluate TargetCall. Table 3 provides details on the evaluated datasets. Datasets D1 & D2, D1 & D4 and D3 & D5 are used for covid detection, sepsis detection and viral detection use cases respectively. For real datasets we randomly sample the datasets provided by prior research to keep a tractable experiment time. We use DeepSimulator to generate the simulated reads [75, 76]. We simulate the dataset D5 due to unavailability of open access raw signal files for viral reads.

**Table 3:** Evaluated read datasets.

| List of Read Datasets | Dataset Type | Number of Reads | Source | DOI Accession |
| --- | --- | --- | --- | --- |
| (D1) Human | Real | 196,000 | [77] | 10.5281/zenodo.7334648 |
| (D2) SARS-CoV-2 | Real | 4,000 | [78] | 10.5281/zenodo.7335539 |
| (D3) Bacterial Mixture (n=7) | Real | 72,567 | [20] | 10.5281/zenodo.7335525 |
| (D4) Bacterial Mixture (n=9) | Real | 15,200 | [20] | 10.5281/zenodo.7335517 |
| (D5) Viral Mixture (n=7) | Simulated | 35,000 | [75, 76] | 10.5281/zenodo.7334592 |

### Evaluated Reference Genomes

We use three reference genomes to evaluate TargetCall on three applications. Table 4 lists the details of these reference genomes.

**Table 4:** List of Reference Genomes Used for each Use Case.

| Reference Genome | Use Case | Datasets Compared | % of Target Reads |
| --- | --- | --- | --- |
| Human (GRCh38) | Sepsis Detection | D1 & D4 | 17.64% |
| SARS-CoV-2 (NC_045512.2) | Covid Detection | D1 & D2 | 2% |
| Viral Combined (n=10) | Viral Detection | D3 & D5 | 32.51% |

For testing viral detection use case, we used 10 viral reference genomes from NCBI RefSeq (with IDs NC_003977.2, AC_000007.1, NC_009334.1, NC_001526.4, NC_010277.2, NC_001731.1, NC_063383.1, NC_055231.1, NC_045512.2, NC_014361.1); and combined them to use as the target reference [79]. Only 7 of these 10 reference genomes are used for simulating reads of dataset D5^1^.

### Evaluation Metrics

We evaluate TargetCall using four different metrics: 1) filtering accuracy, 2) end-to-end accuracy, 3) basecalling execution time, and 4) end-to-end execution time.

We evaluate the filtering accuracy of TargetCall by computing its precision and recall. We define precision as the number of reads that TargetCall correctly labels as on-target, divided by the total number of reads that TargetCall labels as on-target. We define recall as the number of reads that TargetCall correctly labels as on-target, divided by the total number of on-target reads in the dataset. The ground truth on-target reads are determined by the conventional pipeline of basecalling with Bonito and read mapping. An ideal pre-basecalling filter should have 100% recall to maintain accuracy in the downstream analyses and 100% precision to provide maximum performance improvement possible.

For the end-to-end accuracy, we calculate the difference of relative abundances of species after 1) pre-basecalling filtering and 2) conventional basecalling. We compute how much relative abundances (RA) after pre-basecalling filtering is deviated from the true RAs. Equation (1) provides the calculation of the deviation in RAs where *TC_RAi* is the RA of species *i* after TargetCall; and *B*_*RA_i_* is the RA of species *i* after conventional basecalling with Bonito.

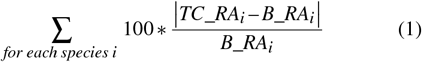

For the basecalling execution time, we compare the time spent on pre-basecalling filtering followed by basecalling of the reads that are accepted by the filter and conventional basecalling. The index generation time of minimap2 is excluded from end-to-end execution time, as this is a one-time task per reference genome.

For the end-to-end execution time, we compare the time spent on entire genome analysis pipeline of basecalling, read mapping and variant calling with and without the use of pre-basecalling filtering.

## 6. Results

### 6.1. Filtering Accuracy

In Figures 3 and 4, we assess the precision and recall of TargetCall with different LightCall configurations for all use cases explained in Section 4, respectively. We make four key observations. First, TargetCall’s precision and recall is between 73.59%-96.03% and 42.57%-99.45% for different configurations of LightCall on average across all three use cases tested, respectively. Second, the precision and recall of TargetCall increases as the model complexity of LightCall increases. Third, increasing the model complexity provides diminishing precision and recall improvements after the *LC_Main_*’s model complexity. Fourth, models smaller than *LC_Main_* are sufficient for use cases with small-to-medium target reference sizes, whereas more complex models are required for use cases with large target reference sizes. We conclude that *LC*_*Main*\*2_ provides the highest precision and highest recall compared to other LightCall configurations.

**Figure 3:**
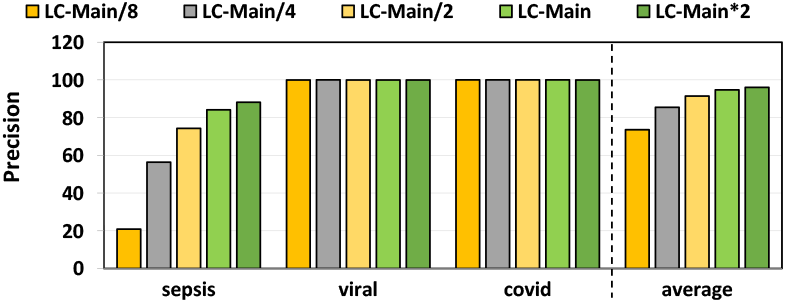
Precision for our evaluated use-cases while using Target-Call with different LightCall configurations.

**Figure 4:**
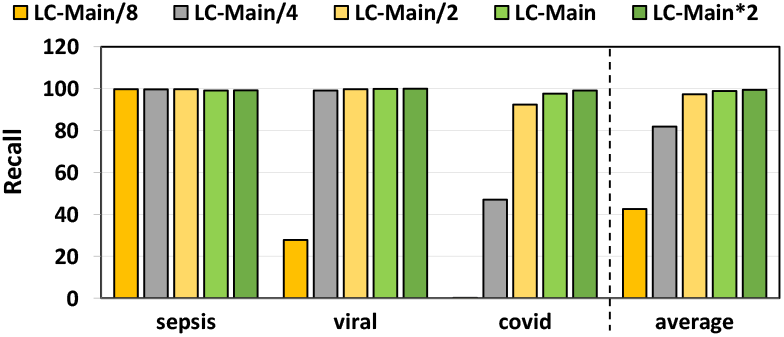
Recall for our evaluated use-cases while using TargetCall with different LightCall configurations.

### 6.2. End-to-End Accuracy

We evaluate the end-to-end accuracy of TargetCall in the viral detection use case. We use Equation (1) to calculate the average deviation in relative abundances as the end-to-end accuracy metric. Ideally, the deviation should be 0%, such that pre-basecalling filtering does not affect the end-to-end accuracy of the genome analysis pipeline at all.

Table 5 lists the average RA deviations calculated for each LightCall configuration. We make the following two key observations. First, the average RA deviations is negligible (≤0.1%) for TargetCall configurations with recall higher than 98.5%. Second, TargetCall’s minor inaccuracy is not biased towards any specific portion of the target reference since otherwise, the deviation of the relative abundances would be higher. Losing a small number of on-target reads randomly enables sequencing depth-of-coverage to compensate for the loss of reads. We conclude that TargetCall’s high sensitivity enables accurate estimates for relative abundance calculations.

**Table 5:** L1 norm for different TargetCall configurations.

| Model Name | Average RA Deviation |
| --- | --- |
| $LC_{Main*2}$ | 0.03% |
| $LC_{Main}$ | 0.08% |
| $LC_{Main/2}$ | 0.23% |
| $LC_{Main/4}$ | 0.91% |
| $LC_{Main/8}$ | 72.19% |

### 6.3. Basecalling Execution Time

Figure 5 provides the total execution time of Bonito and Bonito with TargetCall. We make three key observations. First, TargetCall improves performance of Bonito by 2.13×-3.31×. Second, both precision and performance of LightCall affect the performance of TargetCall, resulting in a non-linear relationship between model complexity and performance. Precision affects the performance of the filter as lower precision results in higher number of falsely accepted reads to be basecalled using conventional basecallers. Model complexity affects the performance of the filter as lower model complexity results in higher LightCall performance with lower precision. Third, decreasing the model complexity increases the performance up to the point where read filtering accuracy is no longer sufficient to filter out reads correctly. This results in a significant number of reads being falsely accepted by the filter, reducing the performance of TargetCall significantly. We conclude that TargetCall significantly improves the execution time of basecalling.

**Figure 5:**
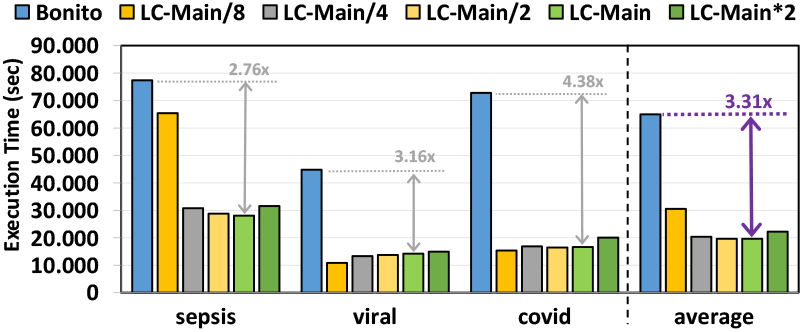
Basecalling execution time for our evaluated use-cases while using TargetCall with different LightCall configurations.

### 6.4. End-to-End Execution Time

Figure 6 provides the total time spent on genome analysis pipeline of basecalling, read mapping and variant calling with and without the use of pre-basecalling filtering. We used minimap2 [56] and DeepVariant [80] for read mapping and variant calling respectively. We make three key observations. First, TargetCall improves the performance of entire genome sequence analysis pipeline by 2.03× -3.00×. Second, the choice of the variant caller affects the end-to-end performance improvement of TargetCall. Since we used a highly-accurate neural network based variant caller, the execution time of variant calling dominated read mapping (not shown). This reduces the end-to-end performance benefits of TargetCall, as the variant calling is performed only on the alignments of on-target reads (i.e., on reduced dataset) determined during relatively light-weight read mapping step. Third, similar to basecalling execution time, multiple factors affect the performance of TargetCall with different LightCall configurations, which results in non-linear relationship between the performance and model complexity of LightCall. We conclude that TargetCall significantly improves the end-to-end execution time of genome sequence analysis pipeline by providing 3.00× speedup.

**Figure 6:**
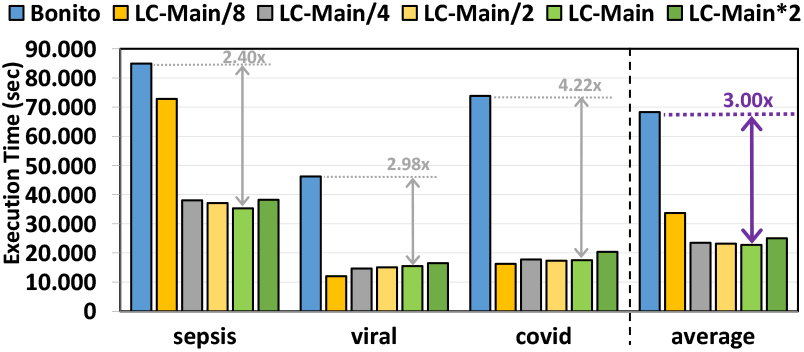
End-to-end execution time for our evaluated use-cases while using TargetCall with different LightCall configurations.

### 6.5. Best Model Selection

We evaluate the performance-accuracy trade-off of TargetCall to determine the best LightCall architecture for TargetCall. Figure 7 aggregates the accuracy (recall) and performance (basecalling speedup) of all LightCall configurations evaluated (except *LC*_*Main*/8_). We make the following three key observations. First, *LC_Main_* provides the highest (3.31×) speedup in basecalling. Second, *LC*_*Main*\*2_ provides the highest (99.45%) accuracy in basecalling. Third, *LC_Main_* provides significantly higher (13.36%) speedup than *LC*_*Main*\*2_ with minimal (0.57%) reductions in accuracy. Therefore, we select *LC_Main_* as the LightCall component of TargetCall. We conclude that TargetCall by using *LC_Main_* improves the performance of basecalling by 3.31 × by precisely filtering 94.71% of the on-target reads while maintaining high (98.88%) sensitivity in filtering.

**Figure 7:**
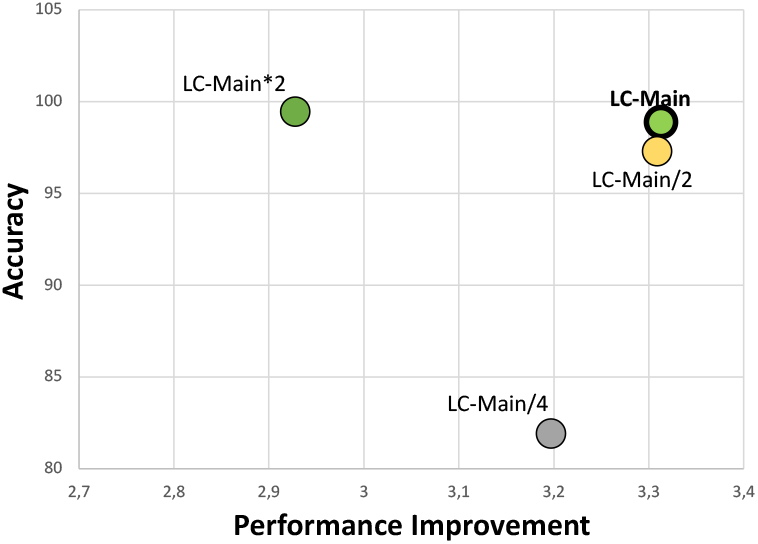
Overall accuracy (recall) and performance (basecalling speedup) improvement for TargetCall with different LightCall configurations.

### 6.6. Comparison to Prior Work

We compare TargetCall with the best LightCall configuration (LC_Main_) with two state-of-the-art targeted sequencing methods UNCALLED and Sigmap. In our evaluations we considered only the two of our use cases where the target reference size is small enough for Sigmap and UNCALLED to be applicable. For the sepsis use case, both tools failed to generate the index structures. We evaluate only these two methods, as they can readily be repurposed as pre-basecalling filters. Sigmap converts the target reference into its *synthetic* raw signal representation, and compares the two raw signals to label the reads as on-target or off-target. UNCALLED probabilistically generates short subsequences of length ~15 from the raw signal, and matches them to the target reference for labelling the reads.

We compare the end-to-end execution time of TargetCall with that of Sigmap and UNCALLED, the results are shown in Figure 8. We observe that TargetCall outperforms Sigmap by 3.07× and UNCALLED by 1.95×. TargetCall’s higher performance benefits come from its higher precision in filtering out off-target reads (not shown).

**Figure 8:**
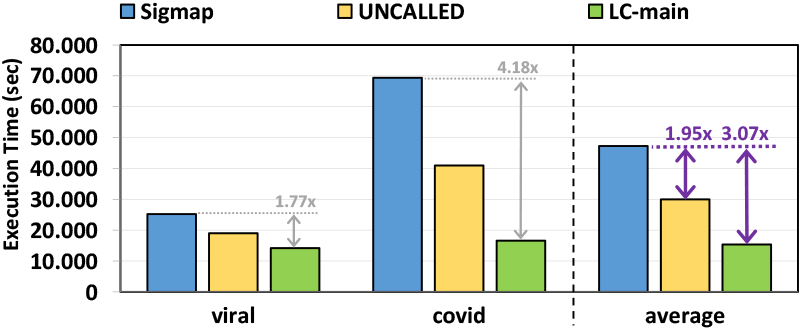
End-to-end execution time of Sigmap, UNCALLED, and TargetCall.

We compare the sensitivity (recall) of TargetCall with that of Sigmap and UNCALLED. The ground truth on-target reads are determined by the conventional pipeline. Figure 9 shows the sensitivity of Sigmap, UNCALLED and TargetCall. We make two key observations. First, TargetCall provides significantly higher sensitivity, +28.3%, than Sigmap on average. Second, TargetCall provides comparable sensitivity to UNCALLED. TargetCall’s sensitivity is higher than UNCALLED for viral detection use case where the target reference is more complex than a single viral reference genome. In contrast, UNCALLED’s sensitivity is higher than TargetCall for the covid detection use case where the target reference is a single small sized viral reference genome. This can be due to UNCALLED’s methodology to be specifically optimized towards small target references, and does not reflect the general trend in other pre-basecalling filtering applications.

**Figure 9:**
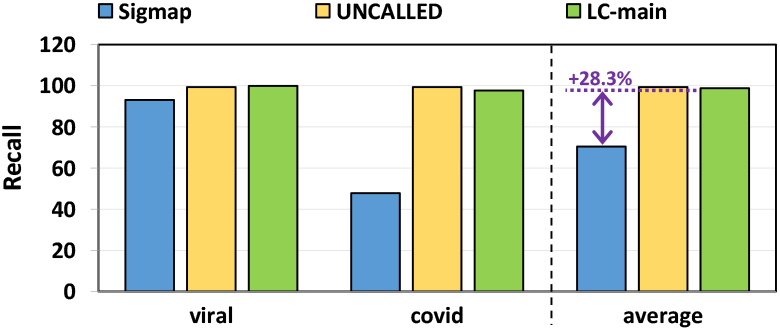
Sensitivity (Recall) of Sigmap, UNCALLED and Target-Call.

We conclude that TargetCall 1) significantly improves the end-to-end execution time of basecalling compared to prior methods by filtering out higher fraction of the off-target reads, 2) is applicable to target reference sizes prior methods are in-applicable, and 3) improves the sensitivity of prior methods on average.

## 7. Related Work

To our knowledge, this is the first work to propose a pre-basecalling filter to eliminate the wasted computation in basecalling. We first discuss previous work in optimizing basecalling. Such methods are orthogonal to our approach, and combining them with pre-basecalling filtering can provide further performance improvements. A complementary set of approaches that discard off-target reads early are called targeted sequencing approaches. Targeted sequencing encapsulates a set of approaches where off-target reads are discarded from the genome analysis pipeline during sequencing [81,82]. We briefly discuss closely-related prior works on targeted sequencing as well as their limitations compared to TargetCall.

### 7.1. Basecalling

Modern basecallers use deep learning-based models that significantly improve the accuracy (≥ 10%) of predicting a nucleotide base from the raw signal compared to traditional non-deep learning-based basecallers [17, 18, 25, 83, 84]. However, these deep learning models use millions of model parameters that makes basecalling computationally extensive. Recent works propose algorithmic optimizations [33, 48] and hardware accelerators [85] to improve the performance of basecallers. These works accelerate the basecalling step without eliminating the wasted computation in basecalling.

### 7.2. Targeted Sequencing

We analyze the targeted sequencing methods in three groups. The first group of work compares the raw electrical signals to a target reference without basecalling the signals in two steps [45]. The first step converts the bases of the reference genome into their *synthetic* raw signal representation. The second step identifies similarities between the raw read signal and the synthetic reference signal. Prior work [45] used the Dynamic Time Warping (DTW) algorithm that measures the similarity of a pair of signals to find the similarity. If the read is not similar to target reference, it is labeled as off-target. Unlike TargetCall, this method is not applicable to large target references of millions of base pairs (bp) long due the quadratic time complexity of DTW with respect to the length of the sequences. To address the computational bottleneck of DTW, Sigmap [19] propose to generate an index for the synthetic reference signal. Sigmap queries the generated index as the reads are being sequenced to find the potential similarity positions between the read and the reference signal, avoiding the DTW calculation. These positions refer to matches of short subsequences of the read and the reference signal, and the final labelling is determined based on clustering these match locations. The use of index structure enables Sigmap to be applicable to target references of length up to ~ 100 Mbp, however, Sigmap is still significantly less applicable compared to TargetCall. Recently, SquigleFilter proposed to design an accelerator to make DTW calculation faster [26] but it was unable to make DTW scalable for large target references.

The second group of works is based on converting the raw signal into a set of bases and comparing these bases to the reference to label the raw signal [28, 46]. Readfish uses a real-time basecalling method to basecall the read as it is being sequenced and perform read mapping on the basecalled portion of the raw signal [46]. Since basecallers are optimized to work on complete reads, this method results in suboptimal base sequences hence may incorrect label the reads [19]. To mitigate these drawbacks, UNCALLED uses an index of the reference genome to probabilistically convert the raw signal into a set of short nucleotide subsequences called seeds and cluster them [28]. The read is classified as on-target read if there is a location in the target reference that has significantly more seeds mapping to that region than the others [28]. UNCALLED is also designed to work *only* on small reference genomes with the goal of performing targeted sequencing [19, 28].

The third group of work uses machine learning to label the raw signals in a sample without performing basecalling and costly analyses in the base space [29, 47]. SquiggleNet [29] can identify a certain class of species with a high accuracy where the class membership is determined based on the target reference. However, this approach requires training the machine learning model for each target reference, which cannot be practically applied to classify any type of species due to the high computational costs of training. Therefore, unlike TargetCall, SquiggleNet cannot be used as a widely-applicable pre-basecalling filter. BaseLess [47] utilizes an array of small neural networks to detect a small subsequences from raw signals and match these subsequences with a target reference that share the same subsequence. This design choice provides a flexible solution that can define the set of pre-trained neural network models of subsequences to identify a certain target reference instead of retraining the neural network model for each species. Unfortunately, none of these works can avoid the cost of training the models multiple times (i.e., for each subsequence or species) to identify the target references in raw signals.

## 8. Conclusion

We propose TargetCall, a pre-basecalling filtering mechanism for eliminating the wasted computation in basecalling. TargetCall performs light-weight basecalling to compute noisy reads using LightCall, and labels these noisy reads as on-target/off-target using Similarity Check. TargetCall eliminates the wasted computation in basecalling by performing basecalling only on the on-target reads. We evaluate TargetCall for three different use cases of pre-basecalling filtering with varying requirements in genome sequence analysis: covid detection, sepsis detection, and viral detection. We show that TargetCall reduces the execution time of basecalling by filtering out majority of the off-target reads, and is more applicable compared to state-of-the-art targeted sequencing methods. We hope that TargetCall inspires future work in pre-basecalling filtering that accelerate other bioinformatics workloads and emerging applications.

## Footnotes

1 Reference Genomes tested can be accessed with DOI 10.5281/zenodo.7335545

